# Antigenic characterisation of human monoclonal antibodies for therapeutic use against H7N9 avian influenza virus

**DOI:** 10.1101/2022.09.16.508351

**Authors:** Pengxiang Chang, Deimante Lukosaityte, Joshua E. Sealy, Pramila Rijal, Jean-Remy Sadeyen, Sushant Bhat, Sylvia Crossley, Rebecca Daines, Kuan-Yin A. Huang, Alain R. Townsend, Munir Iqbal

## Abstract

Since 2013, H7N9 avian influenza viruses (AIVs) have caused more than 1500 human deaths and millions of poultry culling. Despite large-scale poultry vaccination, H7N9 AIVs continue to circulate among poultry in China and pose a threat to human health. Previously, we isolated and generated four monoclonal antibodies (mAbs) derived from humans naturally infected with H7N9 AIV. Here, we investigated the haemagglutinin (HA) epitopes of H7N9 AIV targeted by these mAbs (L3A-44, K9B-122, L4A-14 and L4B-18) using immune escape studies. Our results revealed four key antigenic epitopes at HA amino acid positions 125, 133, 149, and 217. The mutant H7N9 viruses representing escape mutations containing Alanine to Threonine at residue 125 (A125T), Glycine to Glutamic acid at residue 133 (G133E), Asparagine to Aspartic acid at residue 149 (N149D), or Leucine to Glutamine at residue 217 (L217Q) showed reduced or completely abolished cross-reactivity with the mAbs, as measured by hemagglutination inhibition (HI) assay. We further assessed the potential risk of these mutants to humans should they emerge following mAb treatment by measuring the impact of these HA mutations on virus fitness and evasion of host adaptive immunity. Here we showed that the L4A-14 mAb had broad neutralizing capability, and its escape mutant N149D had reduced viral stability and human receptor binding and could be neutralized by both post-infection and antigen-induced sera. Therefore, L4A-14 mAb could be a therapeutic candidate for H7N9 AIV infection in humans and warrants further investigation for therapeutic application.

**IMPORTANCE:** Avian Influenza virus (AIV) H7N9 continues to circulate and evolve in birds, posing a credible threat to humans. Antiviral drugs have been proven useful for the treatment of severe influenza infections in humans, however, concerns have been raised as antiviral resistant mutants have emerged. Monoclonal antibodies (mAbs) have been studied for both prophylactic and therapeutic applications in infectious disease control and have demonstrated great potential. For example, mAb treatment has significantly reduced the risk of people developing severe disease with SARS-COV 2 infection. In addition to the protection efficiency, we should also consider the potential risk of the escape mutants generated by mAb treatment to public health by assessing their viral fitness and potential to compromise host adaptive immunity. Considering these parameters, we assessed four human mAbs derived from humans naturally infected with H7N9 AIV and showed that the mAb L4A-14 displayed potential as a therapeutic candidate.

## INTRODUCTION

Since February 2013, a novel H7N9 avian influenza virus (AIV) has caused 1568 confirmed human infections and 616 deaths, with ~ 40% case fatality rate (1). Though most of the human infections are linked to direct contact with birds or visiting live poultry markets, some hospital or family infection clusters have been observed, raising the concern of possible but limited human-to-human transmission (2). Therefore, H7N9 AIV has been considered a credible pandemic threat.

During early epidemic waves, only low pathogenicity avian influenza (LPAI) virus was detected while the high pathogenicity avian influenza (HPAI) H7N9 virus emerged in late 2016, causing up to 100% mortality in infected chickens (3). Given the threat of H7N9 AIV to human and animal health, the Chinese government implemented a mass vaccination program targeting poultry in 2017. As a result, the number of poultry outbreaks and human infections has dropped dramatically, with only three human infection cases reported during 2016/17 and one human infection case reported during 2017/18; no further human infections have been reported to date (1). However, these viruses have not been eradicated, with continuous sporadic isolation of LPAI and HPAI H7N9 AIV in poultry (4, 5).

Neuraminidase (NA) inhibitors, such as Zanamivir (Relenza) and Oseltamivir (Tamiflu) are the major antivirals that have been recommended for the treatment of severe infection with AIV in humans. However, the rapid emergence of NA inhibitor resistant H7N9 viruses highlights the need for new anti-influenza drugs or therapeutics (6), including the application of human antibodies. Li *et al.* isolated a monoclonal antibody (mAb) (P52E03) recognizing glycine (G) at amino acid residue 133 in the haemagglutinin (HA) and demonstrated its ability to protect against lethal H7N9 AIV challenge in a mouse model (7). Additionally, mAb m826 has been shown to recognize a pH sensitive epitope, also providing full protection of mice challenged with lethal dose of H7N9 AIV (10). Chen *et al.* and Wang *et al.* isolated neutralising antibodies, namely HNIgGD5, HNIgGH8, HNIgGA6 and HNIgGB5, targeting the highly conserved epitopes valine (V) at amino acid residue 186 and leucine (L) at amino acid residue 226 (8, 9) in HA. Another mAb, H7.167, targeting highly conserved epitopes asparagine (N) at amino acid residues 157 and 158, significantly reduces viral lung titre in a mouse intranasal virus challenge study (11).

We previously identified four H7N9 human IgG antibodies, namely, L4A-14, L3A-44, K9B-122 and L4B-18, from humans naturally infected with H7N9 AIV (12). L4-A14 and L3A-14 showed robust therapeutic and prophylactic efficacy against HPAI H7N9 (A/Guangdong/TH005/2017), while L4B-18 failed to protect mice from lethal infection. As for the LPAI H7N9 AIV strain A/Taiwan/1/2013, though L4A-14, L3A-44 and K9B-122 all showed robust prophylactic protection, 100% survival was only found with antibody L4A-14 in a therapeutic model (12).

In this study, we mapped H7N9 HA epitopes using a mAb escape mutant method and investigated the cross-reactivity between these H7HA mAbs with prevalent H7N9 AIVs. The emergence of virus mutants escaping neutralisation following immune pressure, such as that from antibody treatment, is inevitable, therefore it is critical to investigate the risk of these mutants to public health. Efficient human-to-human transmission requires a reduced threshold for pH stability, increased HA thermal stability and the shift of the HA receptor binding affinity from avian to human (13, 14). We therefore assessed the risk of these mAb escape mutants to human health by evaluating the impact of these HA substitutions on receptor binding, pH of fusion, thermal stability, and virus replication fitness. Moreover, we assessed the evasion of vaccine or infection induced humoral adaptive immunity by microneutralization (MN) and haemagglutination inhibition (HI) assays with homologous post-infection or post-vaccination sera.

## Results

### Biological characterization of human mAbs against H7N9 AIV

Four recombinant mAbs, L4A-14, K9B-122, L3A-44 and L4B-18, were produced in ExpiCHO-S™ cells and purified by Protein A affinity chromatography. To assess the neutralizing profile of the mAbs, the HI and MN titres were assessed by using recombinant H7N9 (A/Anhui/2013, referred to as Anhui/13 hereinafter) with HA and NA from H7N9 AIV and the six internal genes from PR8 virus (A/Puerto Rico/8/34 [H1N1]). All four mAbs showed HI activity against H7N9 AIV, with the stronger HI activity observed with L4A-14 followed by L3A-44, while K9B-122 and L4B-18 showed relatively lower HI titre **(Table 1)**. The MN titre followed a similar trend to the HI titre, but no detectable MN activity was found with L4B-18. To investigate whether the mAb can detect denatured HA, the mAbs were used to detect the H7N9 HA by Western blot with denaturing gel electrophoresis. L4A-14 and K9B-122 recognized the H7N9 HA under the denaturing conditions, whereas no H7 HA antigen band was detected with L3A-44 and L4B-18 mAbs **(Figure 1)**.

**FIG 1.**
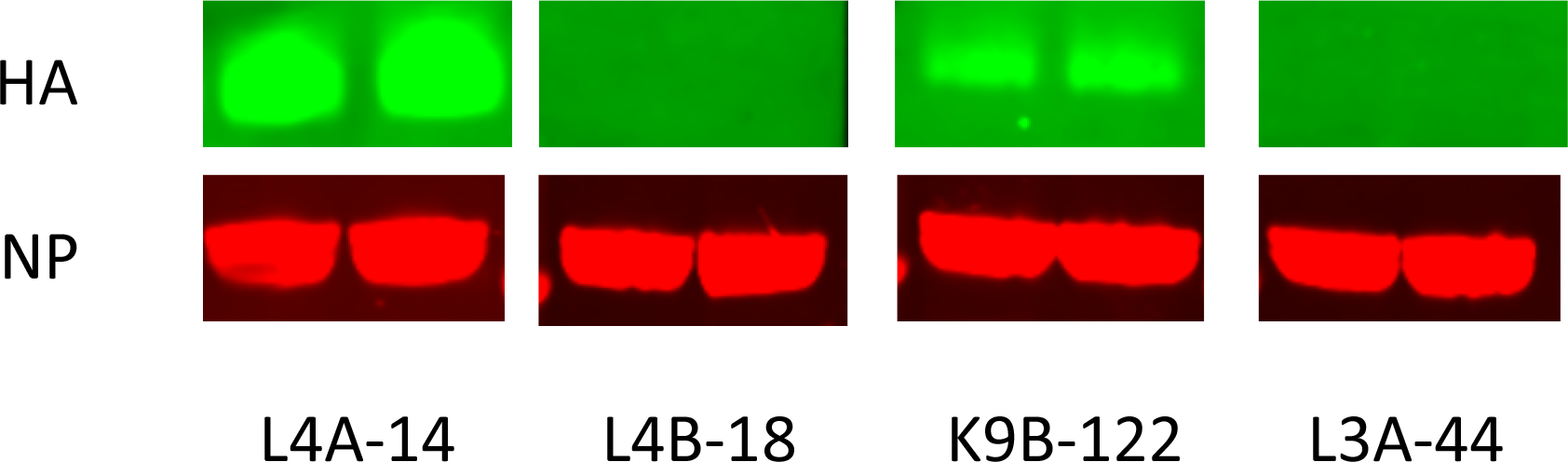
Western blot to detect denatured HA by human anti-H7N9 HA monoclonal antibodies. The HA from H7N9 influenza virus (A/Anhui/1/2013) was probed with human monoclonal antibody L4A-14, L4B-18, K9B-122 or L3A-44 with nucleoprotein (NP) detection as loading control. 3μL of purified H7N9 Anhui/13 viruses (virus concentration of 10 μM) were lysed and loaded per sample with two replicates for each antibody.

**Table 1.**
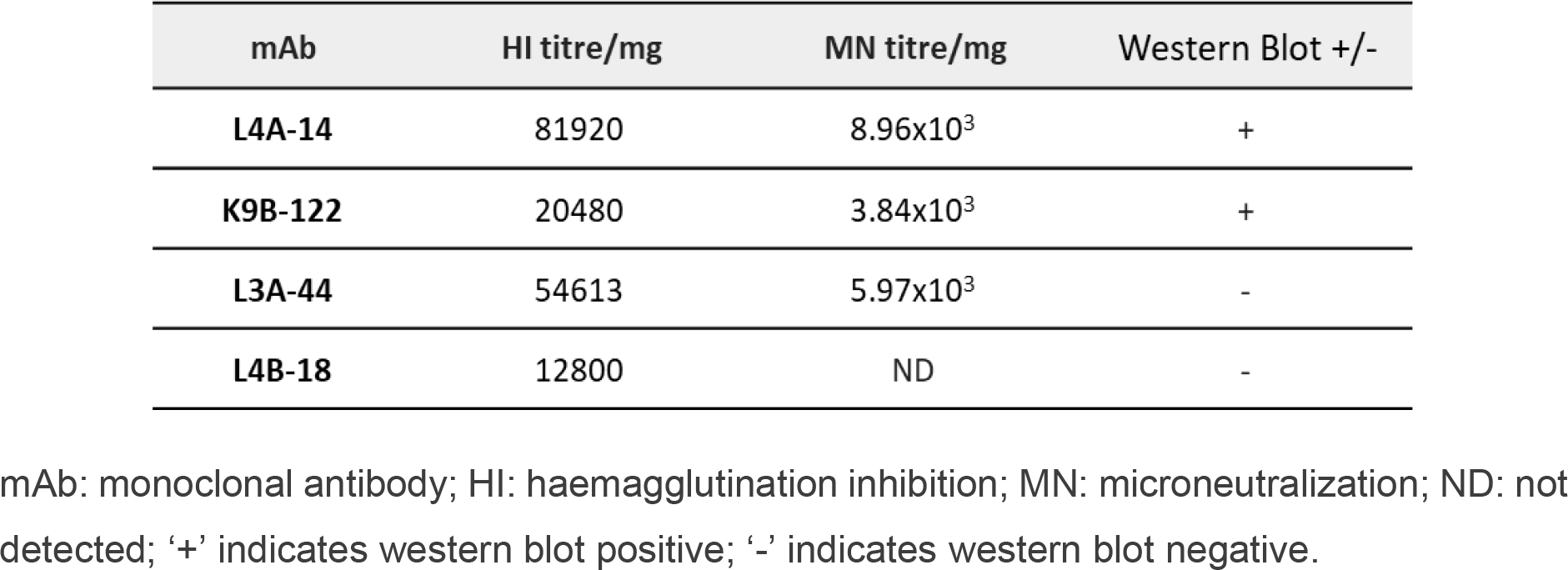
Properties of the human anti-H7N9 monoclonal antibodies.

| mAb | HI titre/mg | MN titre/mg | Western Blot +/- |
| --- | --- | --- | --- |
| L4A-14 | 81920 | 8.96x10<sup>3</sup> | + |
| K9B-122 | 20480 | 3.84x10<sup>3</sup> | + |
| L3A-44 | 54613 | 5.97x10<sup>3</sup> | - |
| L4B-18 | 12800 | ND | - |

mAb: monoclonal antibody; HI: haemagglutination inhibition; MN: microneutralization; ND: not detected; '+' indicates western blot positive; '-' indicates western blot negative.

### H7N9 HA epitope mapping by selection of mAb escape mutants

To select the mAb escape mutants, H7N9 Anhui/13 AIV stock was 10-fold serially diluted and then incubated with 1 mg/mL of each mAb, the antibody/virus mixture was then inoculated into 10-day old embryonated eggs. The HA positive allantoic fluid from the embryonated eggs inoculated with the lowest concentration of virus was selected for viral HA gene sequence analysis.

The results showed that propagation of virus in the presence of the monoclonal antibodies gave rise to the following substitutions: for L4A-14, N to D at residue 149 (N149D); for K9B-122, G to E at residue 133 (G133E); for L3A-44, A to T at residue 125 (A125T); and for L4B-18, L to Q at residue 217 (L217Q) (mature H7 numbering, which will be used throughout; corresponding to H3 numbering N158D, G144E, and A135T and L226Q, respectively). These substitutions are positioned within the respective footprints of the antibodies that selected them (12). In addition, the A125T substitution targeted by L3A-44 resulted in the formation of an N-linked glycan at the amino acid residue 123 (15), which is also within the footprint for antibody L3A-44 (12). N149D is located at the top of the HA, both A125T and L217Q mutations are located within the receptor binding site (RBS), while G133E is located at the edge of the RBS **(Figure 2)**.

**FIG 2.**
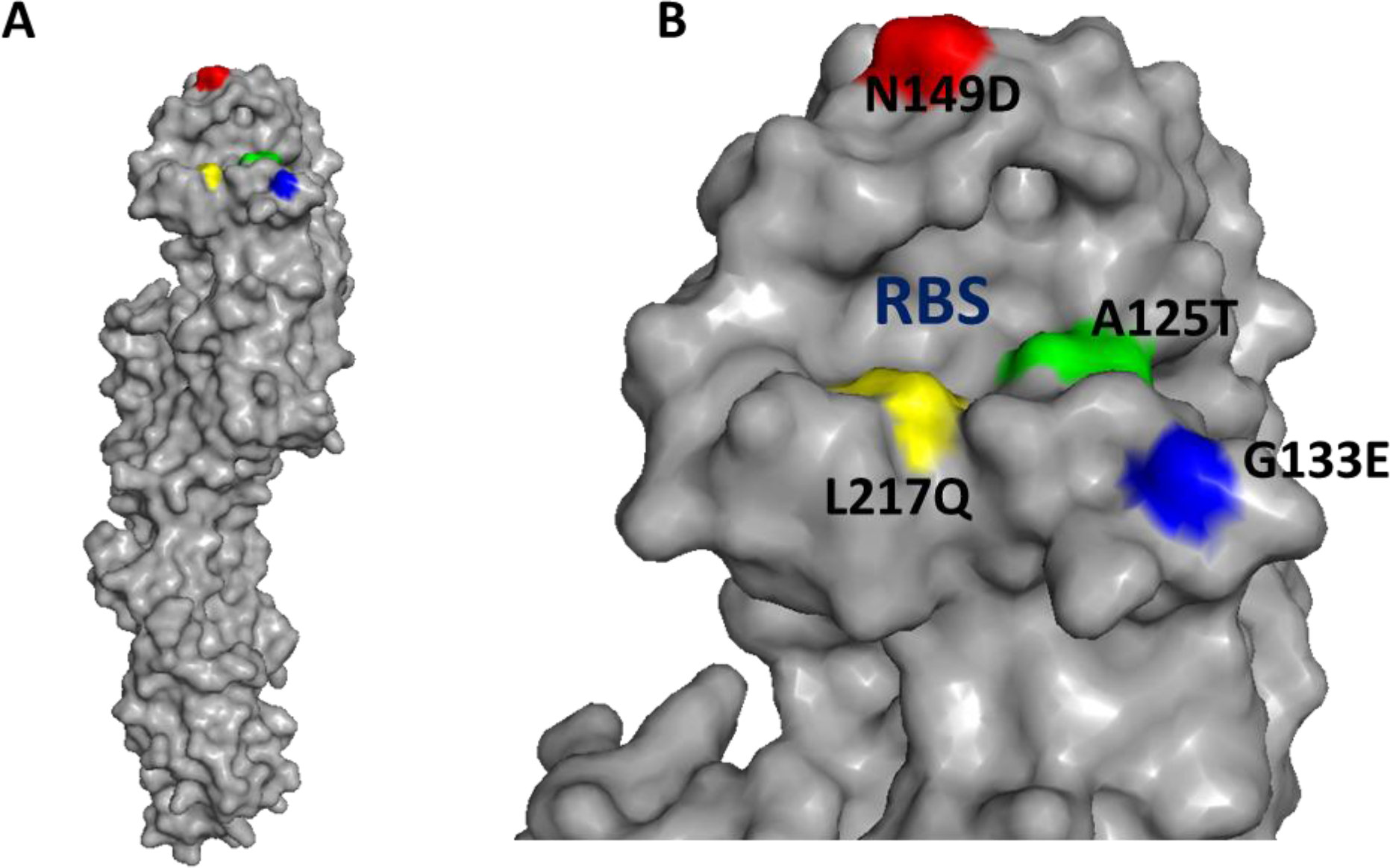
Location of the HA mutation residues on HA monomer (A) and HA1 head domain (B) of three-dimensional structure of H7N9 influenza virus (A/Shanghai/1/2013) (Protein Data Bank [PDB] accession no. 4LN3) (32). RBS, receptor binding site. The mutations A125T, G133E, N149D, and L217Q (mature H7 HA numbering) are indicated in green, blue, red, and yellow respectively.

To confirm a functional role of these substitutions on the escape mutants, each of the identified mutations were engineered into the wild-type HA of Anhui/13 reverse genetic (RG) virus that contained the internal six genes of PR8 virus. The HI assay showed that the mutant virus with HA substitution G133E and A125T completely lost HI activity with mAb K9B-122 and L3A-44 respectively **(Table 2)**. The L217Q mutant virus showed 5 log2 HI titre reduction for L4B-18 mAb while N149D mutant virus showed 2 log2 HI titre reduction with L4A-14 when compared with the parent virus.

**Table 2.** The amino acid substitutions identified in the HA of the human monoclonal antibody escape mutants.

| Selection mAb | Mutations | Reduction in HI titre by escape mutant (log<sub>2</sub>) |
| --- | --- | --- |
| L4A-14 | N149D | 2 (12 → 10) |
| K9B-122 | G133E | 10 (10 → 0) |
| L3A-44 | A125T | 12 (12 → 0) |
| L4B-18 | L217Q | 5 (9 → 4) |

### Cross-reactivity of H7N9 AIV escape mutants with the mAbs

To investigate the cross-reactivity of the H7N9 HA mutants with the individual mAb, all the selected escape mutants were tested by HI assay with the panel of mAbs. Additionally, we included mutant A151T because it introduced an N-linked glycosylation at amino acid residue 149, a regular strategy for influenza viruses to escape the neutralizing antibodies by attaching glycans, which can sterically block the binding of antibodies to multiple antigenic sites (12, 15, 16). We also included the A125V mutant since 95% of the H7N9 AIV isolates in the recent epidemic wave (wave five) contain this mutation in HA (17). The L4A-14 exhibited broad neutralizing activity against almost all the mutants tested, except the A151T substitution in HA which completely abolished its HI activity **(Table 3)**. Interestingly, under the L4A-14 mAb selection pressure, only the N149D mutant emerged rather than the A151T mutant, with only a 2 log_2_ reduction in HI titre. L3A-44 showed reduced cross-reactivity against the G133E mutant, only a slight HI reduction was observed with the A125V and L217Q mutants. L4B-18 showed a moderate reduction in the cross-reactivity with A151T, A125T and A125V mutants, while maintaining its cross-reactivity with N149D and G133E mutants. Among these antibodies, the K9B-122 was most sensitive to mutations, completely losing cross-reactivity with A125V, A125T, L217Q and G133E mutants apart from mutants N149D and A151T.

**Table 3.** Cross-reactivity of H7N9 HA mutants with H7HA mAbs.

| Selection mAb | Mutations | Reduction in HI titre by escape mutant (Log<sub>2</sub>) |  |  |  |
| --- | --- | --- | --- | --- | --- |
|  |  | L4A-14 | L3A-44 | L4B-18 | K9B-122 |
| L4A-14 | N149D | 2 (12 → 10) | 1 (12 → 11) | No change | No change |
|  | A151T | 12 (12 → 0) | 1 (12 → 11) | 2 (9 → 7) | No change |
| L3A-44 | A125T | No change | 12 (12 → 0) | 3 (9 → 6) | 10 (10 → 0) |
|  | A125V | No change | 2 (12 → 10) | 2 (9 → 7) | 10 (10 → 0) |
| L4B-18 | L217Q | 1 (12 → 11) | 2 (12 → 10) | 5 (9 → 4) | 10 (10 → 0) |
| K9B-122 | G133E | No change | 9 (12 → 3) | No change | 10 (10 → 0) |

A151T mutant was additionally included in this analysis because it introduced an N-linked glycosylation at amino acid residue 149 in H7N9 HA; mutant A125V was additionally included since 95% of the H7N9 AIV isolates in recent epidemic wave (wave five) contain this mutation in HA.

### Cross-reactivity of recent H7N9 HA AIV with the mAbs

To investigate the cross-reactivity of the mAbs against recent H7N9 AIV isolates, we performed HI assay against both LPAI (A/Hong Kong/125/2017 [HK125]) and HPAI (A/Guangdong/17SF003/2016 [GDSF003]) from epidemic wave five. Additionally, we included recombinant Anhui/13 with A125T, A151T and L217Q substitutions in HA, as these three mutations have been found in recent H7N9 isolates from poultry in China as well as H7N9 that is serially passaged in the presence of homologous ferret serum (5, 17). The A125T+A151T+L217Q mutant completely lost its cross-reactivity with all the mAbs except for low cross-reactivity to L4B-18 **(Table 4)**. mAb L4A-14 showed strong cross-reactivity with both HPAI GDSF003 and LPAI HK125 while mAb K9B-122 lost cross-reactivity with all the H7N9 AIVs tested apart from Anhui/13, which confirmed earlier results obtained by pseudotype neutralisation (12). The mAbs L3A-44 and L4B-18 still maintained moderate cross-reactivity with HK125 however, a significant reduction in HI titre, 7 log_2_, was observed with GDSF003. These results suggest that mAb L4A-14 harbours robust neutralizing potential with isolates from the epidemic wave five, which presented the greatest rate of human infection with H7N9 AIV.

**Table 4.** Cross-reactivity of mAbs with recent H7N9 AIV.

| mAb<br>(Concentration) | HI titre (Log<sub>2</sub>) |  |  |  |
| --- | --- | --- | --- | --- |
|  | Anhui/13 | GDSF003 | HK125 | A125T+A151T+L217Q |
| L4A-14 (1.0 mg/ml) | 12 | 10 | 12 | 0 |
| L3A-44 (1.5 mg/ml) | 12 | 3 | 8 | 0 |
| L4B-18 (0.8 mg/ml) | 9 | 2 | 7 | 1 |
| K9B-122 (1.0 mg/ml) | 10 | 0 | 0 | 0 |

Anhui/13: recombinant H7N9 A/Anhui/2013; HK125: recombinant A/Hong Kong/125/2017; GDSF003: recombinant H7N9 A/Guangdong/17SF003/2016; A125T+A151T+L217Q: recombinant Anhui/13 with HA mutations A125T, A151T and L217Q.

### Cross-reactivity of H7N9 AIV escape mutants with the homologous sera

It is inevitable that escape mutants will emerge following the therapeutic use of mAbs. Therefore, it is critical to assess the risk of these mutants to public health before their application as a therapeutic for human H7N9 AIV infection. Ideally, the escape mutant should not be able to evade infection or vaccine induced immunity. To this end, we checked the HI and MN titre of the mAb escape mutants with both chicken and ferret post-infection sera. Additionally, we included the chicken antiserum raised against H7N9 inactivated virus to mock chicken post-vaccination serum. In agreement with a previous study, L217Q resulted in a significant drop in HI titre to both ferret post-infection serum (4 log_2_) and chicken post-vaccination serum (3 log_2_) (**Table 5**) (17). However, only 1 log_2_ change in HI titre were observed with chicken post-infection serum. The A125T and A151T substitutions in HA resulted in a slight reduction in HI titre to both ferret post-infection serum and chicken post-vaccination serum, whereas no change was observed with chicken post-infection serum. The G133E mutant showed comparable HI titre with wild-type to both chicken post-infection and post-vaccination sera. Interestingly, a 1 log_2_ increase in HI titre was observed with ferret post-infection serum. The N149D substitution in HA had no obvious impact on the HI titre against chicken vaccination serum, but surprisingly a 1 log_2_ increase in titre was observed with both ferret and chicken post-infection serum. Consistent with the HI assay, all the sera tested showed reduced MN titre against L217Q while no significant changes were observed with N149D and A151T mutant viruses **(Figure 3A, 3B and 3C).** In contrast to the HI assay results, the ferret and chicken post-infection serum showed a significant reduction in MN titre against A125T and G133E mutants, while no significant change was observed with chicken post-vaccination serum. To conclude, the A125T, G133E and L217Q substitution in HA resulted in significant decrease in cross-reactivity to both chicken and ferret post-infection serum, however, only L217Q mutant showed reduced cross-reactivity against chicken post-vaccination serum when compared to the wild-type Anhui/13. All the post-infection and post-vaccination sera tested here demonstrated comparable neutralizing profiles among wild-type Anhui/13, mAb L4A-14 escape mutant N149D and mutant A151T, having completely lost HI activity against mAb L4A-14.

**Table 5.** Cross-reactivity of H7N9 AIV escape mutants with the homologous sera.

| Mutation | HI titre (Log<sub>2</sub>) |  |  |
| --- | --- | --- | --- |
|  | Ferret serum (Post-infection) | Chicken serum (Post-Infection) | Chicken serum (Post-Vaccination) |
| Wild-type | 10 | 8.3 | 9 |
| A125T | 8 | 8.3 | 8 |
| G133E | 11 | 8.3 | 9 |
| N149D | 11 | 9.3 | 9 |
| A151T | 9 | 8.3 | 8 |
| L217Q | 6 | 7.3 | 6 |

**FIG 3.**
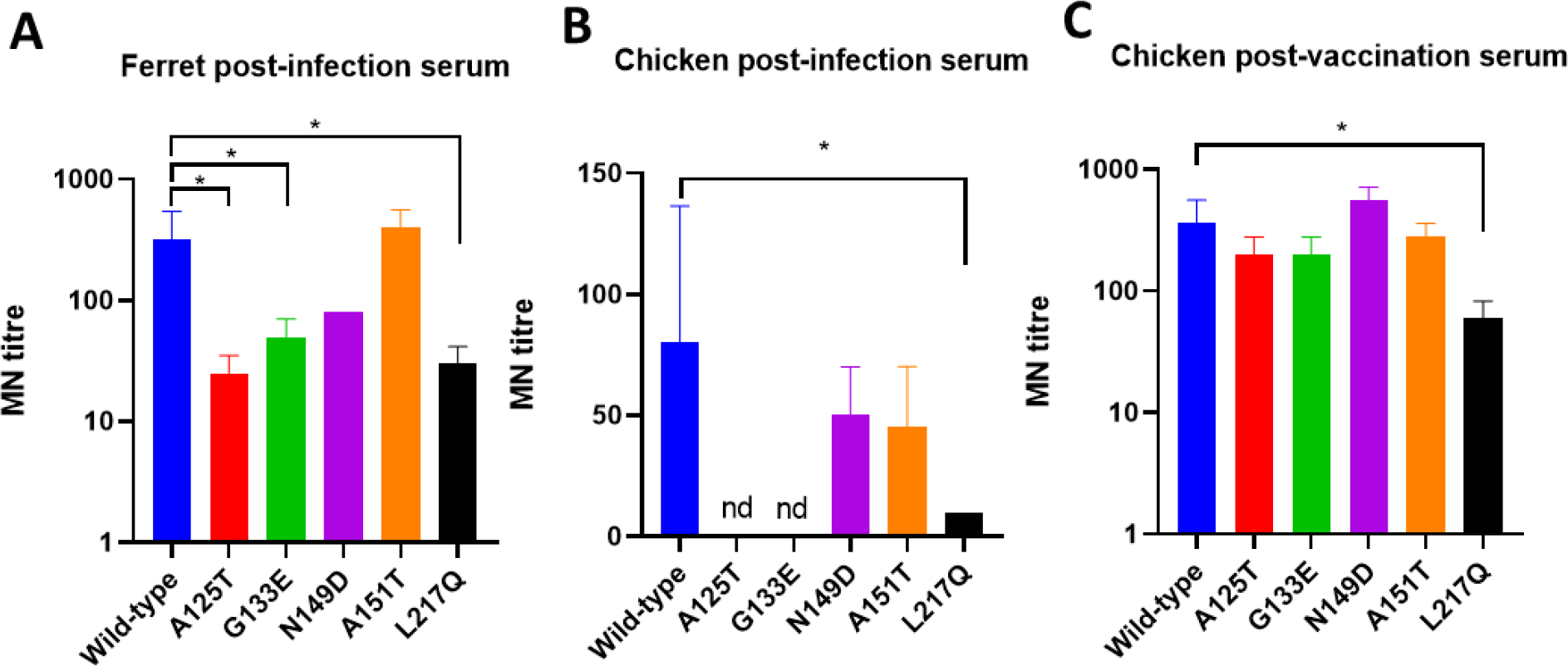
Neutralization by ferret post-infection serum (A), chicken post-infection serum (B) and chicken post-vaccination serum (C) against Anhui/13 wild-type and its mAb escape mutants. A151T mutant was included since it resulted in glycosylation at amino acid residue 149 and completely lost cross-reactivity with mAb L4A-14. Error bar = standard error of mean. * p < 0.05; nd = no detection. MN = microneutralization.

### A125T and N149D mutants replicated robustly *in vitro* and *in ovo*

To evaluate the effects of mAb escape mutations in the HA on virus replication fitness, we generated a series of reverse genetics-based mutant viruses carrying the escape mutations A125T, G133E, N149D and L217Q by site-directed mutagenesis. We did not include A151T mutant for further analysis because it did not emerge under mAb pressure. Moreover, the A151T substitution in HA had minimal impact on the virus neutralizing profile against homologous sera, presented a significant drop in viral thermal stability and reduced receptor binding were observed previously so is therefore unlikely to pose increased zoonotic risk (15).

We assessed the propagation for each of the viruses in mammalian Madin-Darby canine kidney (MDCK) cells and MDCK-SIAT1 (SIAT) cells, which have been modified to express a higher density of α2,6 sialyltransferase (18). Similarly to the wild-type, the L217Q mutant replicated poorly in MDCK cells, despite higher titres at 15 hours post-infection (h.p.i) **(Figure 4A)**. The A125T, G133E and N149D mutants replicated significantly better than the wild-type in MDCK cells at all the time points except for 24 h.p.i, where only mutant N149D replicated to a significant higher titre than the wild-type **(Figure 4A)**. As for the SIAT cells, both A125T and N149D mutants replicated to a significant higher titre than the wild-type at 24, 48 and 72 h.p.i, with only N149D significant higher than wild-type at 15 h.p.i **(Figure 4B)**. Both L217Q and G133E mutants replicated comparably to wild-type, though the titre for the G133E mutant is slightly higher than the wild-type throughout. To further assess the effects of these mutations on virus replication, 10-day old embryonated eggs were inoculated with 100 plaque-forming units of Anhui/13 wild-type and its mutants for 48 hours, with virus titres determined by HA assay **(Figure 4C)** and plaque assay **(Figure 4D)**. The replication of L217Q mutant was comparable to Anhui/13 wild-type while other all mutants showed a significant higher replication titre. The plaque assay results showed similar pattern to the HA assay, except that the replication titre for the G133E mutant was not statistically different than the wild-type Anhui/13 virus **(Figure 4C/4D)**. To conclude, the mAb escape mutants, A125T and N149D mutants demonstrated enhanced replication fitness both *in vitro* and *in ovo*.

**FIG 4.**
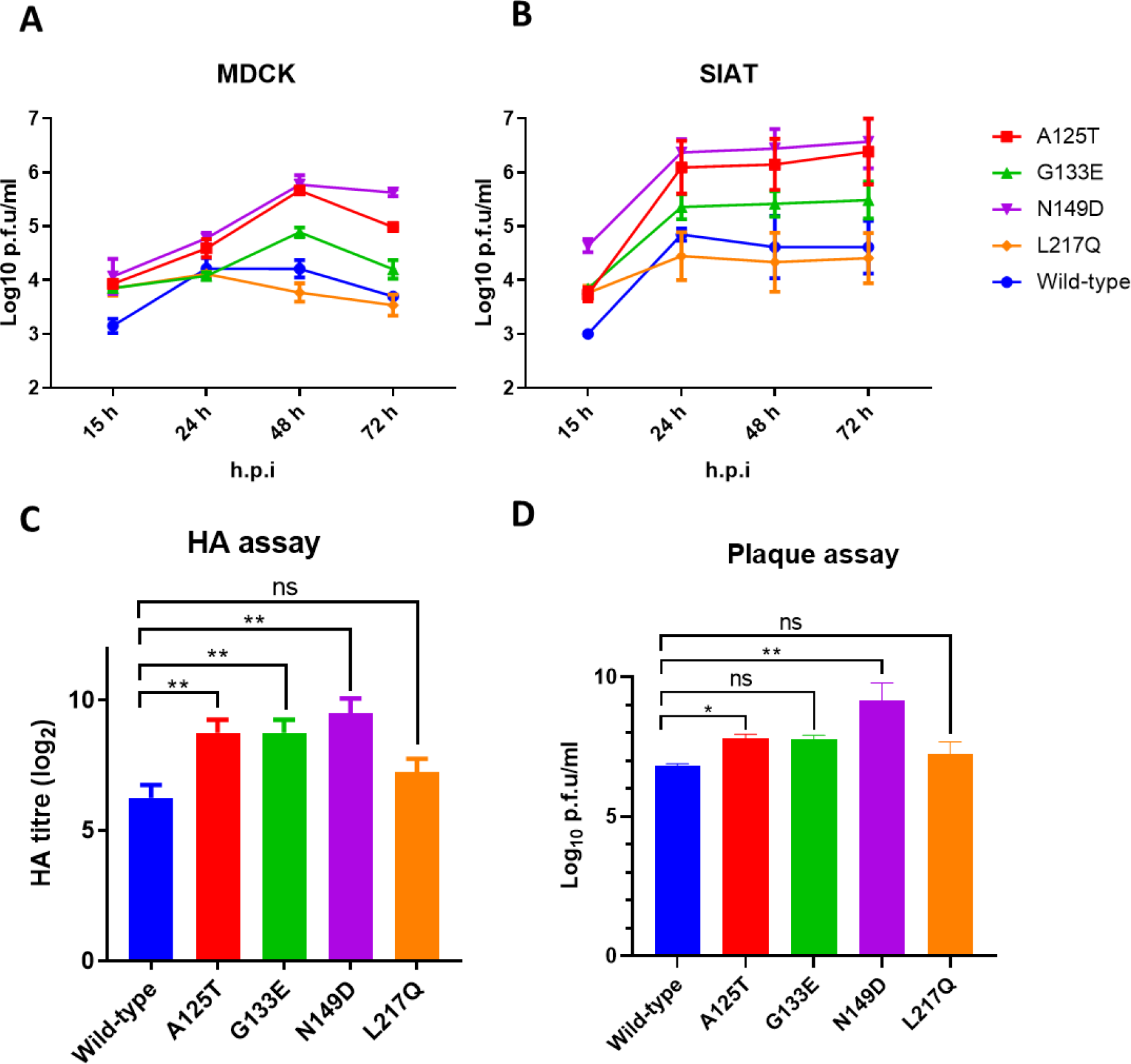
Replication of recombinant Anhui/13 and its mAb escape mutants *in ovo*, MDCK and SIAT cells. The replication kinetics of Anhui/13 wild-type and its mutants in **(A)** MDCK and **(B)** SIAT cells (MDCK cells modified to express a higher density of α2,6 human receptors). Cells were infected with a multiplicity of infection (MOI) of 0.001, supernatants were collected at the 15−, 24−, 48- and 72-hour post-infection, and the virus titres were determined by plaque assays in MDCK cells (statistical analysis in Supplementary Table 1 and 2). 10-day old embryonated eggs were inoculated with 100 plaque-forming units of Anhui/13 wild-type and its mutants for 48 hours before allantoic fluid was harvested and the viral replication determined by **(C)** HA assay and **(D)** plaque assay. Error bar = standard error of mean. * p < 0.05; ** p < 0.001; ns = not significant.

### mAb escape mutations affected H7N9 AIV receptor-binding properties

Receptor binding affinity is one of the key determinants for the interspecies transmission of influenza viruses. To examine the impacts of mAb escape mutations on virus receptor binding properties, we utilized biolayer interferometry to characterize the receptor-binding profiles of these viruses to the avian-like and human-like receptor analogues, 3′-sialylacetyllactosamine (3SLN) and 6′-sialylacetyllactosamine (6SLN). In agreement with previous reports, the H7N9 Anhui/13 showed comparable binding to both 3SLN and 6SLN receptor analogues, and the L217Q mutant showed a considerable increase in binding towards avian 3SLN (~432 folds), with only a slight decrease (~2 folds) to 6SLN analogues **(Figure 5)** (15, 19). Substitutions A125T, G133E and N149D reduced the receptor binding towards both 3SLN and 6SLN analogues. Compared with the A125T mutant (~2-fold reduction to 3SLN and ~4-fold reduction to 6SLN), there was slightly greater reduction with mutant G133E (~7-fold reduction to 3SLN and ~4-fold reduction to 6SLN) and mutant N149D (~9-fold reduction to 3SLN and ~7-fold reduction to 6SLN).

**FIG 5.**
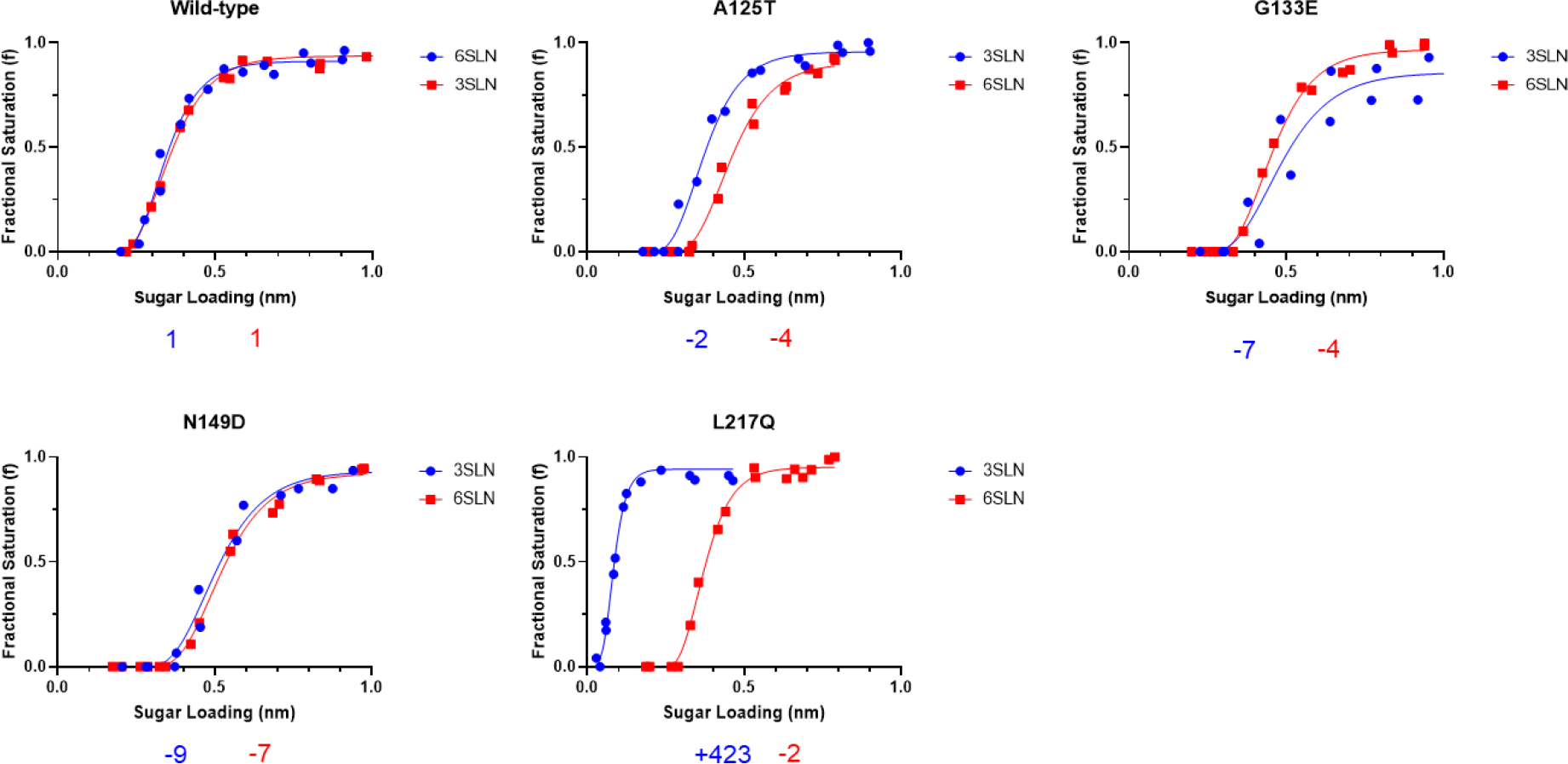
Receptor binding profiles of recombinant Anhui/13 and its mAb escape mutants. The binding of purified recombinant virus Anhui/13 (wild-type) and A125T, G133E, N149D and L217Q mutants to avian (α2,3-SLN, shown in blue) and human (α2,6-SLN, shown in red) receptor analogues by biolayer interferometry. The numbers below each figure show the fold change of receptor binding of indicated viruses as compared to recombinant Anhui/13 (wild-type). ‘−’ indicates reduction, ‘+’ indicates increase. Data is the combination of two repeats for each virus and receptor analogue combination.

### mAb escape mutations affected H7N9 AIV pH fusion and thermal stability

The pH of fusion has been shown to play an important role in host adaptation and transmission (20). Consistent with the previous report, Anhui/13 wild-type triggered fusion at pH 5.6 or lower **(Figure 6A)** (21). The mAb escape mutant A125T, G133E and N149D showed a similar pH fusion threshold as the Anhui/13 wild-type. Only the L217Q substitution in HA caused a slight decrease in pH fusion threshold.

**FIG 6.**
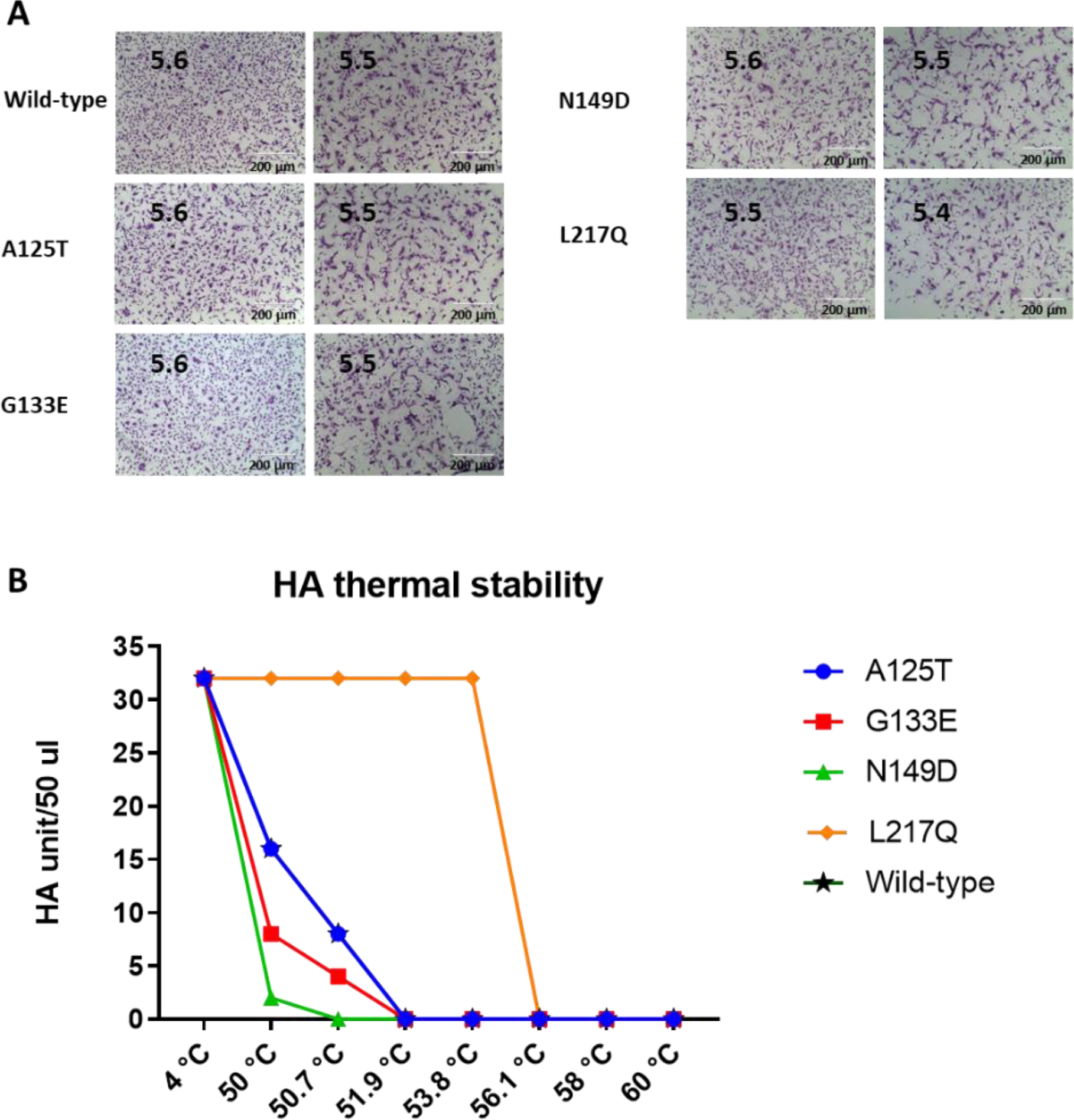
pH fusion and thermal stability of recombinant Anhui/13 and its mAb escape mutants. **(A)** Syncytium formation in Vero cells infected with recombinant Anhui/13 (wild-type) and A125T, G133E, N149D and L217Q mutants. The pH, at which 50% of maximum syncytium formation was observed, was taken as the predicted pH of fusion, shown on the left panel. The syncytium formation at 0.1 pH unit lower than the fusion threshold were shown on the right panel as control. Results shown are representative of three experimental repeats. **(B)** HA thermal stability of recombinant Anhui/13 wild-type and A125T, G133E, N149D and L217Q mutants, with the Anhui/13 wild-type as control. 32 haemagglutinating units (HAU) of recombinant virus were either left at 4°C fridge as control or heated at 50°C, 50.7°C, 51.9°C, 53.8°C, 56.1°C, 58.0°C, 59.2°C and 60°C for 30 min before the HA assay. Results shown are representative of three experimental repeats.

In addition to receptor binding and pH of fusion, HA thermal stability also plays a vital role in AIV evolution (13, 14). Anhui/13 wild-type and the mAb escape mutants were heated at 50°C, 50.7°C, 51.9°C, 53.8°C, 56.1°C, 58.0°C, 59.2°C and 60°C for 30 min, after which the loss of HA activity was determined by HA assay. The G133E and A125T mutants showed a similar heat stability profile as the Anhui/13 wild-type. The L217Q mutation dramatically increased the HA stability, while the N149D slightly reduced stability **(Figure 6B)**.

## DISCUSSION

In our previous study, we have isolated four potent human monoclonal antibodies from humans naturally infected with H7N9 AIV. To gain in depth knowledge of the therapeutic potential of these antibodies, we mapped the antigenic epitopes of these antibodies by escape mutant selection methods. In addition to the neutralizing capability, we also evaluated the potential risk to humans of mAb-induced escape mutants by assessing their replication fitness, receptor binding, pH fusion, thermal stability, and potential to evade host adaptive immunity. Overall, the L4A-14 mAb showed robust neutralizing capability and broad cross-reactivity with H7N9 AIVs. Moreover, its escape mutant N149D demonstrated reduced thermal stability while retaining strong antigenic cross-reactivity with both post-infection and vaccination sera, making L4A-14 mAb a suitable therapeutic candidate for further investigation.

H7N9 AIV is still circulating among poultry in China, frequently causing human infection. Fortunately, few cases of human infection have occurred since the Chinese government introduced large-scale poultry vaccinations. We previously used an *in vitro* immune escape mutant selection method to model H7N9 evolution under immune pressure (17). We predicted that the mutations A125T, A151T and L217Q in HA might occur under the pressure of humoral immunity, all of which have been found in recent strains of H7N9 AIV isolated from poultry in China (5). Here, we found that A125T+A151T+L217Q mutant lost the cross-reactivity with almost all mAbs, maintaining only limited cross-reactivity with L4B-18 mAb, implying that these mAbs may provide little protection against H7N9 AIV containing such mutations. However, it is noteworthy that these mutations, particularly A125T and A151T double mutations in HA, resulted in the complete loss of human receptor binding and decreased HA thermal stability, making them less likely to emerge in a H7N9 AIV pandemic (15). This is further supported by the finding that only one human infection case of H7N9 AIV contains both A125T and A151T mutations in the HA (22). Though A125T and A151T mutations resulted in the complete loss of cross-reactivity with L3A-44 and L4A-14 mAbs respectively, the L4A-14 mAb retained antigenic cross-reactivity against L3A-44 mAb escape mutants and *vice versa*. Therefore, antibody treatment containing a combination of L3A-44 and L4A-14 mAbs could be a highly efficient therapeutic combination in neutralizing containing A125T or A151T substitution.

Interestingly, L4A-14 induced the escape mutant N149D rather than the A151T mutant, with significant antigenic change. One possible explanation is that A151T substitutions result in considerable reductions in HA stability and receptor binding, negatively impacting viral fitness, compared to mutant N149D (15).

The Q217L substitution in HA has been linked to a binding affinity switch of avian receptors to human receptors (α2,3 to α2,6) in H2, H3 and H4 avian influenza viruses (23–25), which could be a prerequisite for zoonotic infection. The L217Q substitution has been shown to hinder transmission of H7N9 AIV between pigs (26). Similarly, Sun *et al*., reported that LPAI H7N9 containing L217 transmitted well between ferrets while two HPAI H7N9 viruses containing Q217 could not (27). Here, we showed that L217Q mutants emerged under the L4B-18 mAb pressure *in ovo*. Further investigation is required to justify application of this mAb for passive immunization *in vivo*, with the aim to drive the virus mutation L217Q to lower pandemic potential.

We previously showed that the amino acid at position 217 is a key mediator of H7N9 AIV antigenicity. Both HI and MN results support this finding, demonstrating that the L217Q substitution in HA resulted in significant reduction in antigenic cross-reactivity with both ferret and chicken post-infection serum, as well as chicken post-vaccination serum. In contrast, Wang *et al.* concluded that L217Q had minimal impact on the antigenicity of H7N9 AIV (28) however, this discrepancy could be due to the different model animals used, mouse and macaque, and we agree that multiple serological assays should be taken into consideration when evaluating antigenic variation. Despite no HI titre change was observed with the A125T mutant, a considerable reduction in MN titre was detected with chicken post-infection serum. Similarly, despite comparable HI titre, G133E mutant showed significant decrease in MN titre to both chicken and ferret post-infection serum. Interestingly, we found that A125T, G133E and L217Q caused significant antigenic change by both post-infection serum of chicken and ferret. However, only substitution L217Q resulted in significant antigenic change to the post-vaccination serum of both hosts. It appeared that the post-vaccination serum from chickens vaccinated with adjuvanted inactivated virus is more broadly cross-reactive than the post-infection serum. This difference could be due to the effects of adjuvant, which has been shown to affect virus antigenicity (29). Unfortunately, the chicken post-vaccination and post-virus infection sera were not produced in the same experiment with proper control, and thus warrants further investigation to understand the conflicting phenomenon found in this study.

To conclude, we evaluated the potential of utilizing human-derived mAbs raised against H7N9 AIV as an infection therapy. However, immune pressure, including mAb therapy will inevitably induce escape mutations. As such, we identified and mapped the H7N9 HA by immune escape *in ovo*. Additionally, we assessed the potential zoonotic risk and the viral fitness consequence of the identified mutations. Our data revealed that L4A-14 mAb could be a potential therapeutic candidate for human H7N9 infections, with escape mutants posing a lower risk to human health and therefore pandemic potential.

## MATERIALS AND METHODS

### Ethics Statement

The embryonated chicken egg work was carried out in accordance with the guidance and regulations of United Kingdom Home Office regulations under project licence number P68D44CF4. All the influenza virus related work was carried out in biosafety level two conditions.

### Viruses and cells

The recombinant H7N9 viruses containing HA and neuraminidase (NA) from low pathogenic avian influenza (LPAI) H7N9 (A/Anhui/1/2013) (referred to as wild-type Anhui/13) and the internal gene segments from A/Puerto Rico/8/34 (H1N1) (PR8) were generated as previously described (17). The recombinant H7N9 AIV A125T, A151T and L217Q mutants of Anhui/13 were generated via site-directed mutagenesis with primers described in previous study (17). The primers for site-directed mutagenesis of G133E and N149D are listed in **Table 6**. Rescued viruses were propagated in 10-day-old embryonated chicken eggs and virus stocks were kept at −80°C.

**Table 6.**
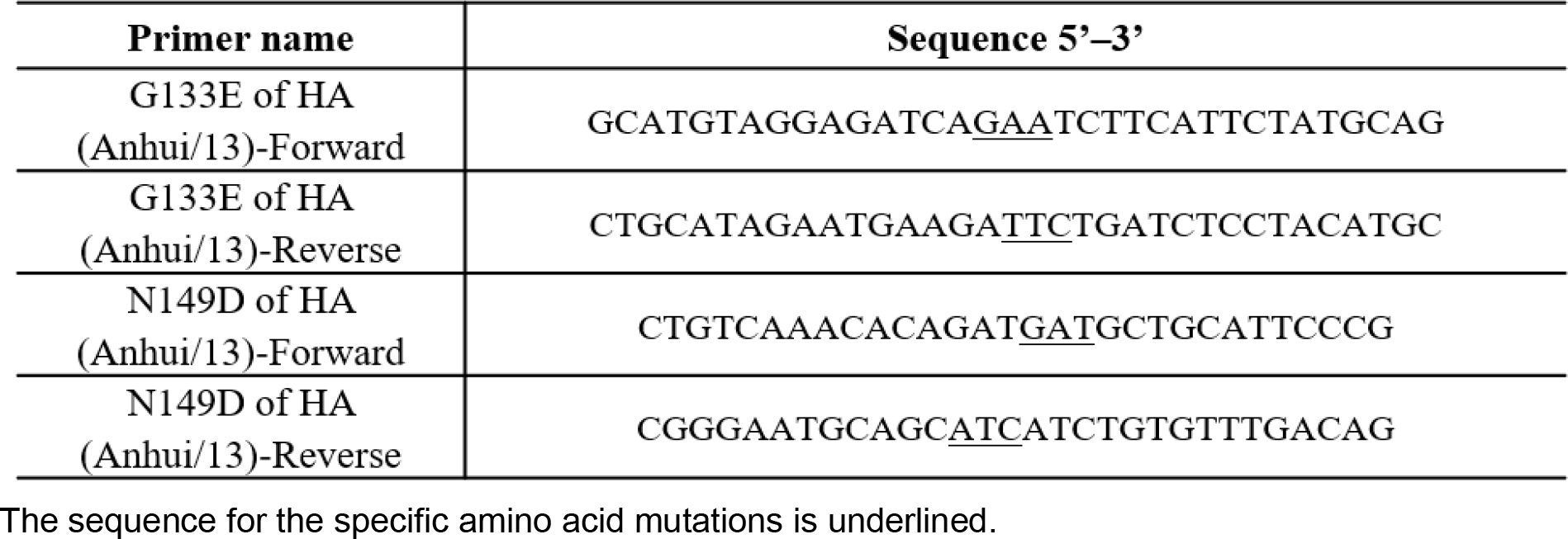
List of primers for H7N9 HA amplification and site-specific mutations.

The Madin-Darby canine kidney (MDCK), MDCK-SIAT1 (SIAT) cells, human embryonic kidney (HEK) 293T and Vero cells (ATCC) were maintained with Dulbecco’s Modified Eagle’s medium (DMEM) (Gibco), supplemented with 10% fetal calf sera (FCS) (Gibco), 100 U/mL Penicillin-Streptomycin (Gibco) at 37°C under a 5% CO_2_ atmosphere.

### Monoclonal antibodies and sera

The monoclonal antibodies were produced in ExpiCHO-S™ cells and purified by protein affinity chromatography. Basically, the ExpiCHO-S™ cells were co-transfected with antibody light and heavy chain plasmids using ExpiFectamine™ CHO Transfection Kit according to manufacturer’s protocol. The supernatants were filtered and purified by HiTrap Protein G HP antibody purification columns (GE Healthcare).

Post-infection ferret antiserum raised against LPAI H7N9 (A/Anhui/1/2013) virus was kindly provided by John McCauley (The Francis Crick Institute, UK). The Post-infection chicken antiserum was raised previously in our lab by infection of SPF-derived Rhode Island red chickens (Roslin Institute, UK) with recombinant LPAI H7N9 (A/Anhui/1/2013) virus (30). The sera were heat-inactivated at 56°C for 30 min before further analysis.

### Replication kinetics in MDCK and MDCK-SIAT1 cells

MDCK or MDCK-SIAT1 cells were infected with H7N9 AIV at multiplicity of infection (MOI) 0.001 and incubated at 37 °C for 1 hour (h). The cells were then washed once with phosphate-buffered saline (PBS) (central services unit at The Pirbright Institute) and replenished with DMEM containing 2 μg/ml tosylsulfonyl phenylalanyl chloromethyl ketone (TPCK)-treated trypsin. The supernatants were taken at 15, 24, 48 and 72 h post-infection and stored in −80 °C before titration by plaque assay in MDCK cells. The plaque assay was carried out as previously described (31). MDCK cells in 12-well plate were infected with H7N9 AIV for 1 h before the infection medium was removed. Cells were then washed once with PBS before addition of agarose overlay. The cells were further incubated at 37°C for 72 h before being stained with 1% crystal violet (Sigma-Aldrich).

### *In ovo* growth

Ten-day-old embryonated chicken eggs were infected with 100 plaque-forming units (pfu) of indicated recombinant H7N9 AIV. The eggs were incubated at 35°C for 48 h before allantoic fluid was harvested for hemagglutination (HA) assay and plaque assay.

### HA assay

The HA titration assays were performed using 1% chicken red blood cells (RBC) as described in the WHO animal influenza training manual (32). Briefly, 50 μl of virus was two-fold serially diluted and added to the 96-well V- bottom shaped plates (Greiner), followed by addition of 50 μl of 1% chicken RBCs. The plates were incubated for 30 minutes (min) at room temperature before the HA titre was recorded and presented as HA units/50 μl.

### HA thermal stability assay

The recombinant H7N9 AIV were diluted with embryonated chicken eggs allantoic fluid to 32 HA units/50 μl. The viruses were then left at 4°C as control or heated at 50°C, 50.7°C, 51.9°C, 53.8°C, 56.1°C, 58.0°C, 59.2°C and 60°C using PCR thermal cycler (Biorad) for 30 min before HA assay.

### Syncytium formation assays

The pH of fusion for H7N9 AIV was determined by syncytium formation assays as previously described (33). Briefly, the Vero cells in 96-well plate were infected with H7N9 AIV, the inoculum was aspirated at 1 h post-infection and washed once with PBS before addition of DMEM with 10% FCS. At 16 h post-infection, cells were washed once with PBS and treated with DMEM medium containing 3 μg/ml TPCK-treated trypsin for 15 min. The cells were then exposed to PBS buffer with pH values ranging from 5.2 to 6.0 (at 0.1 unit increments) for 5 min. The PBS buffer was then replaced with DMEM with 10% FCS and the cells were further incubated for 3 h at 37°C before being fixed with methanol and acetone (1:1 in volume) mixture and stained with 20% Giemsa stain (Sigma-Aldrich) for 1 h at room temperature. The pH at which 50% of maximum syncytium formation estimated was taken as the predicted pH of fusion. Images were taken on the Evos XL cell imaging system (Life Technologies).

### Virus Microneutralisation (MN) Assay

Monoclonal antibody (1 mg/ml) or polyclonal antisera were two-fold serially diluted and then mixed with an equal volume of 100 pfu recombinant H7N9 AIV in serum-free medium. MDCK cells were washed once with PBS before infection with the virus-serum or virus-antibody mixture for 1 h at 37°C, the cells were then washed once with PBS and replenished with DMEM containing 2 μg/ml TPCK-treated trypsin. The cells were further kept at 37°C for 72 h before crystal violet staining.

### Western blot

Three microliters of the sucrose-gradient-purified viruses (virus concentration of 10 μM, which was determined using an enzyme-linked immunosorbent assay against the virus NP as described previously [28]) were mixed with NuPAGE LDS sample buffer (4X) and NuPAGE™ Sample Reducing Agent (10X), and heated at 70°C for 10 min before being loaded onto NuPAGE 4 to 12% bis-Tris protein gels (Life Technologies). The proteins were transferred using iBlot™ 2 Transfer Stacks (Life Technologies) and the membrane was blocked with 5% dried skimmed milk (Marvel). The HA and NP protein were probed with human anti-H7N9 HA monoclonal antibodies and mouse anti-NP monoclonal antibody (ATCC) respectively followed by incubation with IRDye680RD goat anti-mouse IgG and IRDye800CW goat anti-human IgG secondary antibody and visualized using Odyssey Clx (LI-COR).

### Biolayer interferometry

The influenza virus receptor binding affinity was measured on an Octet RED instrument (ForteBio). The equilibrium responses for virus binding were plotted as a function of the amount of sugar immobilized on the biosensor calculated from the response during the sugar loading step (34). Briefly, influenza viruses were purified by continuous 30 to 60% (w/v) sucrose gradient and the concentration of viruses was determined using an enzyme-linked immunosorbent assay against the virus NP as described previously (34). The Biotinylated α2,3- and α2,6-linked sialyl lactosamine sugars (3SLN and 6SLN, respectively) were purchased from GlycoNZ. Virus was diluted in HBS-EP buffer (TEKnova) containing 10 μM oseltamivir carboxylate (Roche) and 10 μM zanamivir (GSK) to the final concentration of 100 pM. Equilibrium responses for virus binding were plotted as a function of the amount of sugar immobilized on the biosensor calculated from the response during the sugar loading step (34).

### Statistical analysis

Statistical analysis was performed using GraphPad Prism 8 (GraphPad Software). One-way ANOVA was used to test differences between different groups for figure 3A, 3B, 3C, 4C and 4D. The two-way ANOVA analysis (Tukey’s multiple comparisons test) was used for figure 4A and 4B. p values < 0.05 were considered significant.

## ACKNOWLEDGMENTS

The work described herein was funded by the UK Research and Innovation (UKRI), Biotechnology and Biological Sciences Research Council (BBSRC) grants: BB/T013087/1, UK-China-Philippines-Thailand Swine and Poultry Research Initiative (BB/R012679/1), Zoonoses and Emerging Livestock Systems (ZELS) (BB/L018853/1 and BB/S013792/1), the Global Challenges Research Fund (GCRF) One Health Poultry Hub (BB/S011269/1), the Responsive Mode (BB/N002571/1), Japan Partnering Award (BB/P016472/1), the Pirbright Institute strategic program grants (BBS/E/I/00007030, BBS/E/I/00007031, BBS/E/I/00007035 and BBS/E/I/00007036), the British Council Newton Fund Institutional Links grant (IL3261727271) and the Royal Society Newton Fund grant (NAF\R1\191166). The funders had no role in study design, data collection, data interpretation or the decision to submit the work for publication.

We would like to thank Steve Martin for the authorization to use Octet analysis software for the receptor binding analysis.

